# *Pseudomonas aeruginosa* biofilms display carbohydrate ligands for CD206 and CD209 that interfere with their receptor function

**DOI:** 10.1101/2020.04.20.051292

**Authors:** Sonali Singh, Yasir Almuhanna, Mohammad Y. Alshahrani, Douglas Lowman, Peter J. Rice, Chris Gell, Zuchao Ma, Bridget Graves, Darryl Jackson, Kelly Lee, Rucha Kelkar, Janice Koranteng, Dan Mitchell, Ana da Silva, Farah Hussain, Gokhan Yilmaz, Francesca Mastrotto, Yasuhiko Irie, Paul Williams, David Williams, Miguel Camara, Luisa Martinez-Pomares

## Abstract

Bacterial biofilms represent a challenge to the healthcare system because of their resilience against antimicrobials and immune attack. Biofilms consist of bacterial aggregates embedded in an extracellular polymeric substance (EPS) composed of carbohydrate polymers, nucleic acids and proteins. Carbohydrates within *P. aeruginosa* biofilms include neutral and mannose-rich Psl, and cationic Pel composed of *N*-acetyl-galactosamine and *N*-acetyl-glucosamine. Here we show that *P. aeruginosa* biofilms display ligands for the C-type lectin receptors mannose receptor (MR, CD206) and Dendritic Cell-Specific Intercellular adhesion molecule-3-Grabbing Non-integrin (DC-SIGN, CD209). Binding of MR and DC-SIGN to *P. aeruginosa* biofilms is carbohydrate-and calcium-dependent and extends to biofilms formed by clinical isolates. Confocal analysis of *P. aeruginosa* biofilms shows abundant DC-SIGN ligands among bacteria aggregates while MR ligands concentrate into discrete clusters. DC-SIGN ligands are also detected in planktonic *P. aeruginosa* cultures and depend on the presence of the common polysaccharide antigen. Carbohydrates purified from *P. aeruginosa* biofilms are recognised by DC-SIGN and MR; both receptors preferentially bind the high molecular weight fraction (HMW; >132,000Da) with K_D_s in the nM range. HMW preparations contain 74.9-80.9% mannose, display α-mannan segments and alter the morphology of human dendritic cells without causing obvious changes in cytokine responses. Finally, HMW interferes with the endocytic activity of cell-associated MR and DC-SIGN. This work identifies MR and DC-SIGN as receptors for bacterial biofilms and highlights the potential for biofilm-associated carbohydrates as immunomodulators through engagement of C-type lectin receptors.

**Author Summary:** Selective engagement of pattern recognition receptors during infection guides the decision-making process during induction of immune responses. This work identifies mannose-rich carbohydrates within bacterial biofilms as novel molecular patterns associated with bacterial infections. *P. aeruginosa* biofilms and biofilm-derived carbohydrates bind two important lectin receptors, MR (CD206) and DC-SIGN (CD209), involved in recognition of self and immune evasion. Abundance of MR and DC-SIGN ligands in the context of *P. aeruginosa* biofilms could impact immune responses and promote chronic infection.

## Introduction

*Pseudomonas aeruginosa* is a versatile opportunistic pathogen that causes acute infection after invasive procedures and burns, and chronic infections in patients with persistent lung disease and compromised immunity (1). *P. aeruginosa* infection is especially troublesome in people with cystic fibrosis where it is a major determinant of irreversible loss of lung function and mortality (2, 3). Niches created by hospital procedures such as the use of catheters and implants as well as contact lenses are effectively colonised by *P. aeruginosa* which exploits an armoury of cell-associated and secreted virulence determinants that facilitate invasion and establishment of infection (4). Transition from planktonic to sessile growth and biofilm development are central to *P. aeruginosa* pathogenesis (1, 4, 5). Biofilms contribute to *P. aeruginosa* persistence by increasing tolerance to anti-microbial agents and immune defences (1). Within such bacterial communities, the cells are embedded within an extracellular polymeric substance (EPS) matrix composed of carbohydrates, nucleic acids and proteins (6, 7). *P. aeruginosa* produces three major carbohydrates: Psl, Pel and alginate, with Psl and Pel playing major roles in biofilm formation in a strain-dependent manner (6–8). Psl is neutral and mannose-rich (9). Pel is cationic and largely composed of *N*-acetyl-galactosamine and *N*-acetyl-glucosamine (10). Here we tested the hypothesis that *P. aeruginosa* biofilms could directly engage lectin receptors expressed by immune cells. In particular, the high mannose content of Psl suggested potential binding to mannose-binding C-type lectin receptors (CLRs) such as mannose receptor (MR, CD206) (11) and Dendritic Cell-Specific Intercellular adhesion molecule-3-Grabbing Non-integrin (DC-SIGN, CD209) that are predominantly expressed by selected populations of macrophages and dendritic cells (MR and DC-SIGN) and non-vascular endothelium (MR) (11, 12). The roles ascribed to these molecules are numerous and include promotion of antigen presentation and modulation of cellular activation (11, 12). MR contains two independent carbohydrate-binding domains, the cysteine-rich domain (MR-CR) and the C-type lectin-like domains (MR-CTLD4-7) that recognise sulfated and mannosylated sugars, respectively (11). DC-SIGN binds to high mannose structures and blood type Lewis antigens through its extracellular region (12). Here we demonstrate that MR and DC-SIGN ligands are present within *P. aeruginosa* biofilms. Distinct binding pattern of both lectins highlights the heterogeneity of carbohydrate structures within the biofilm structure. In addition, DC-SIGN recognises ligands in planktonic *P. aeruginosa* cultures that depend on the presence of the common polysaccharide antigen (13). Carbohydrates purified from biofilm cultures, particularly high molecular weight species, bind MR and DC-SIGN and interfere with their endocytic activity. These results demonstrate the capacity of *P. aeruginosa* biofilm-associated carbohydrates to engage immune receptors and suggest an active role for these structures in modulating the immune responses to biofilms.

## Results

### *P. aeruginosa* biofilms display DC-SIGN and MR ligands

The mannose-rich nature of carbohydrates produced by *P. aeruginosa* biofilms (9) suggested the possibility of immune mannose-specific lectins recognising these structures. We tested whether the lectins MR and DC-SIGN bound *P. aeruginosa* biofilms by analysing the interaction of recombinant Fc chimeric molecules DC-SIGN-Fc and MR-CTLD4-7-Fc (14) to biofilms generated in 96 well plates as described in Materials and Methods. Both DC-SIGN and MR-CTLD4-7 bound to *P. aeruginosa* PAO1 biofilms and DC-SIGN displayed higher binding compared to MR (Figure 1A). Inhibition assays using selected monosaccharides, confirmed that MR and DC-SIGN binding to PAO1 biofilms is carbohydrate-dependent and preferentially inhibited by mannose and fucose, in agreement with the sugar specificity previously shown for both lectins (Figure 1A), and Ca2+-dependent (Figure 1B), as expected for these C-type lectins (11, 12). Recognition by MR and DC-SIGN was not restricted to biofilms generated by PAO1 and extends to biofilms generated by wound clinical isolates (CW 2, CW3, CW5-B, CW5-S, CW6 and CW7). This strain collection comprised isolates collected from bone (CW2 and CW5-B), wound tissue (CW3, CW6 and CW7) and blood (CW5-S) and two serotypes (13)(CW2, CW3, CW5 and CW6 are serotype O6) and CW7 is part of the serotype O5 cluster, which includes O5, O18, and O20 serotypes. Sequence analysis confirmed presence of the *psl* operon (8) in all isolates although point mutations could alter the levels and structure of the carbohydrate (Figure S1). DC-SIGN and MR-CTLD4-7 binding to biofilms formed by clinical isolates follows a similar pattern to that found for PAO1, i.e. increased binding by DC-SIGN compared to MR (Figure 1C).

**Figure 1.**
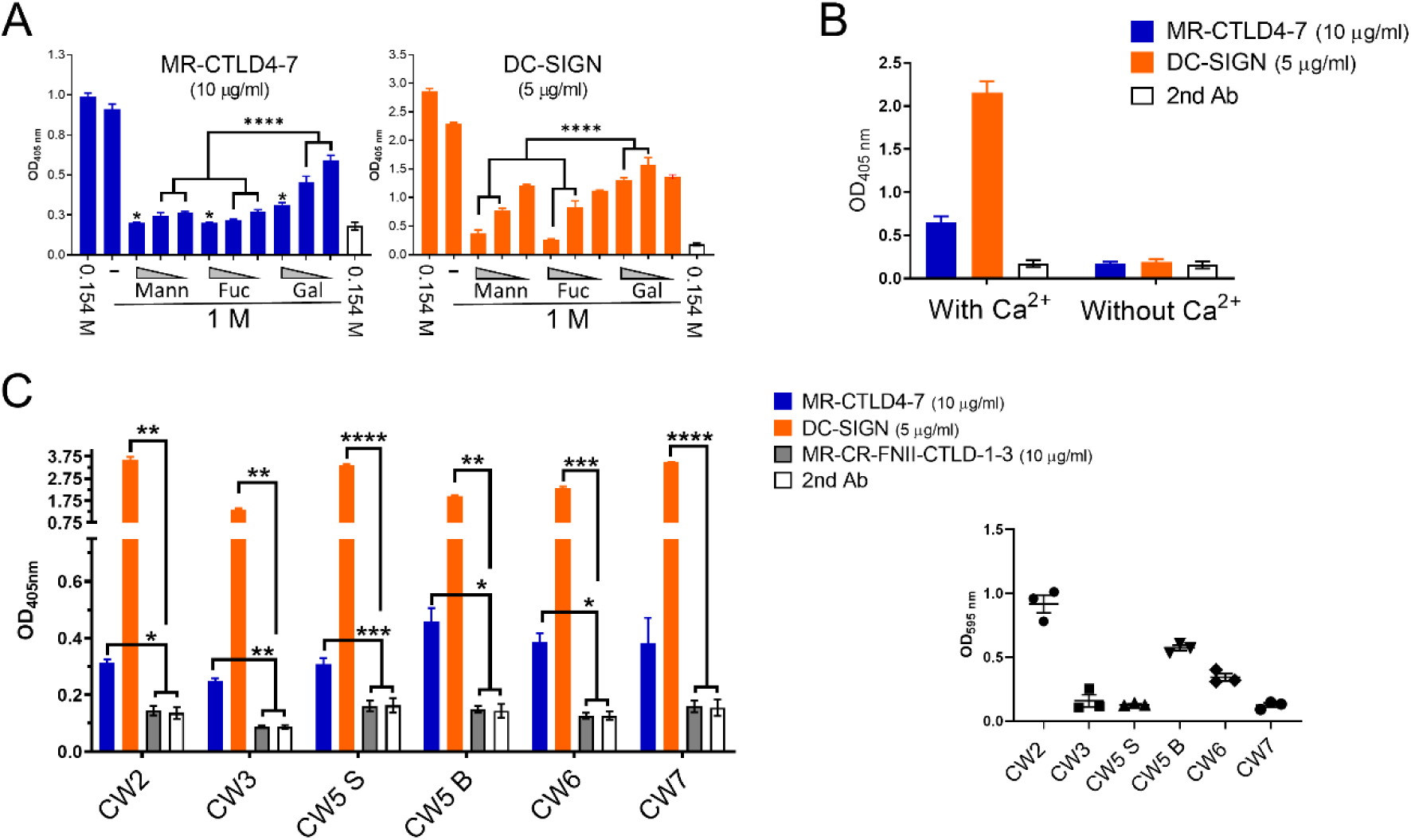
*P. aeruginosa* biofilms display ligands for MR and DC-SIGN. *P. aeruginosa* biofilms generated in 96 well plates for 24 h were fixed and incubated with MR-CTLD4-7-Fc or DC-SIGN-Fc followed by anti-human Fc antibody conjugated to alkaline phosphatase. **A.** DC-SIGN and MR-CTLD4-7 bind to PAO1 biofilms and binding is selectively inhibited by mannose and fucose compared to galactose. Binding assays were performed in two NaCl concentrations (0.154 M and 1M) and for the 1M condition, incubation with Fc chimeric proteins was performed in the presence or absence of different concentrations of monosaccharides (1, 0.2 and 0.04 mM). Binding at 1M NaCl is comparable to that observed at 0.154 M (ns). Presence of monosaccharides significantly reduced protein binding. Mannose and fucose preferentially inhibited binding of MR-CTLD4-7 compared to galactose at 0.2 and 0.04 mM and binding of DC-SIGN at 1 and 0.2 mM. Mann: mannose, Fuc: fucose; Gal: galactose. N=3 in triplicate. One-way ANOVA with Tukey’s multiple comparison test. **B.** DC-SIGN and MR-CTLD4-7 binding to PAO1 biofilms is Ca^2+^-dependent. Binding of MR-CTLD4-7 and DC-SIGN to PAO1 biofilms was tested in the presence and absence of 10 mM CaCl_2_. N=2 in triplicate. **C.** MR and DC-SIGN recognise biofilms formed by *P. aeruginosa* wound isolates. Binding of both lectins was significant in all instances except in the case of MR-CTLD4-7 binding to CW7. No binding of the control protein MR-CR-CR-FNII-CTLD1-3 (15) was observed. Two-way ANOVA with Tukey’s multiple comparison test. Right panel: Biofilm formation by clinical isolates tested using crystal violet assay. N=3 in triplicate. Graphs show mean +/− SEM. ns: non-significant.

### Distinct distribution of DC-SIGN and MR ligands within PAO1 biofilms

Confocal analysis of DC-SIGN and MR-CTLD4-7 ligands within PAO1 biofilms generated under flow conditions (Figure S2) unveiled unique ligand distribution for both lectins (Figure 2). In accordance with the strong binding detected using ELISA-based assays (Figure 1) DC-SIGN ligands within biofilms were widely distributed and particularly abundant between bacteria aggregates with some areas displaying substantial ligand density (Figure 2 top panel; Video S1). MR-CTLD4-7 ligands within *P. aeruginosa* biofilms were less abundant and displayed a granular distribution forming clusters located both among and on bacteria aggregates (Figure 2, bottom panel; Video S2).

**Figure 2.**
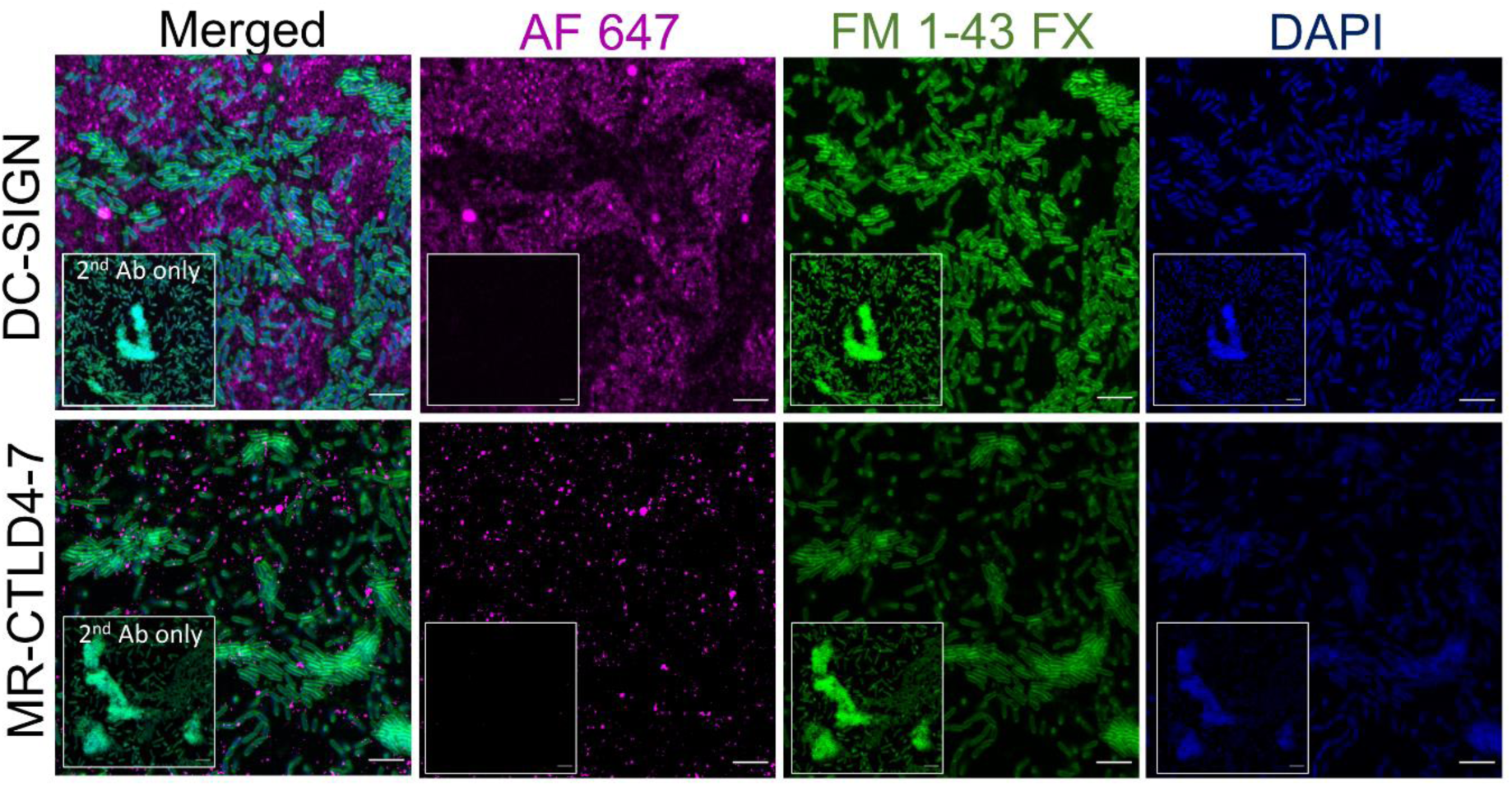
Distribution of MR and DC-SIGN ligands in *P. aeruginosa* biofilms. PAO1 biofilms generated under flow conditions for 18h as described in Figure *S2* were incubated with DC-SIGN and MR-CTLD4-7 Fc-chimeric proteins followed by anti-human Fc-secondary antibody conjugated to Alexa 647 (magenta) and counterstained with DAPI (DNA, blue) and FM1-43FX (bacteria, green). Z-stacks were acquired for all samples using confocal microscopy (see videos S1 and S2). The top panels show slice 9 for DC-SIGN, and 12 for corresponding secondary Ab control (insets) and bottom panels slice 12 for MR-CTLD4-7 and its control (insets). The same settings for image acquisition and processing were maintained for test and control samples (Videos S3 and S4). Inset shows signal with biofilms incubated with secondary antibody only. No specific labelling was seen with MR-CR-FNII-CTLD1-3 Fc chimeric protein (video S5). Size bar 4 µm.

### DC-SIGN binds to planktonic *P. aeruginosa*

Further analysis of DC-SIGN and MR binding to *P. aeruginosa* biofilms using ELISA-based assays identified binding of DC-SIGN to biofilms generated by the Psl-deficient mutant Δ*wsp*F Δ*psl* (16)(Table 1); this mutant is not expected to contain mannose-rich carbohydrates. The Δ*wspF* background confers constitutive high levels of cyclic-di-GMP, overproduces Psl and promotes biofilm formation; a phenotype that resembles that of small rough colony variants found during chronic infection (17). MR-CTLD4-7 displays reduced binding to Δ*wsp*F Δ*psl* biofilms indicating that the *psl* operon is likely responsible for the generation of MR-CTLD4-7 ligands (Figure 3A). We investigated the possibility of DC-SIGN interacting with planktonic cells, which could account for binding to biofilms in the absence of Psl. DC-SIGN binds planktonic *P. aeruginosa* cultures and binding was independent of Psl and/or Pel (Figure 3B). MR-CTLD4-7 did not bind planktonic PAO1. *P. aeruginosa* produces two forms of O antigen; a homopolymer of D-rhamnose trisaccharide repeats named common polysaccharide antigen (CPA) or A band, and a heteropolymer that consists of repeating units of three to five distinct sugars named as O-specific antigen (OSA) or B band (13). Deletions in *wbpM*, *rmd* or *wbpL* in PAO1 causes loss of CPA, OSA, or both, respectively (18). Binding of planktonic *P. aeruginosa* to DC-SIGN required expression of *rmd* or *wbpL* (Figure 3C) suggesting the requirement for CPA LPS and a broader role for DC-SIGN, compared to MR, in the recognition of *P. aeruginosa* infection.

**Table 1.**
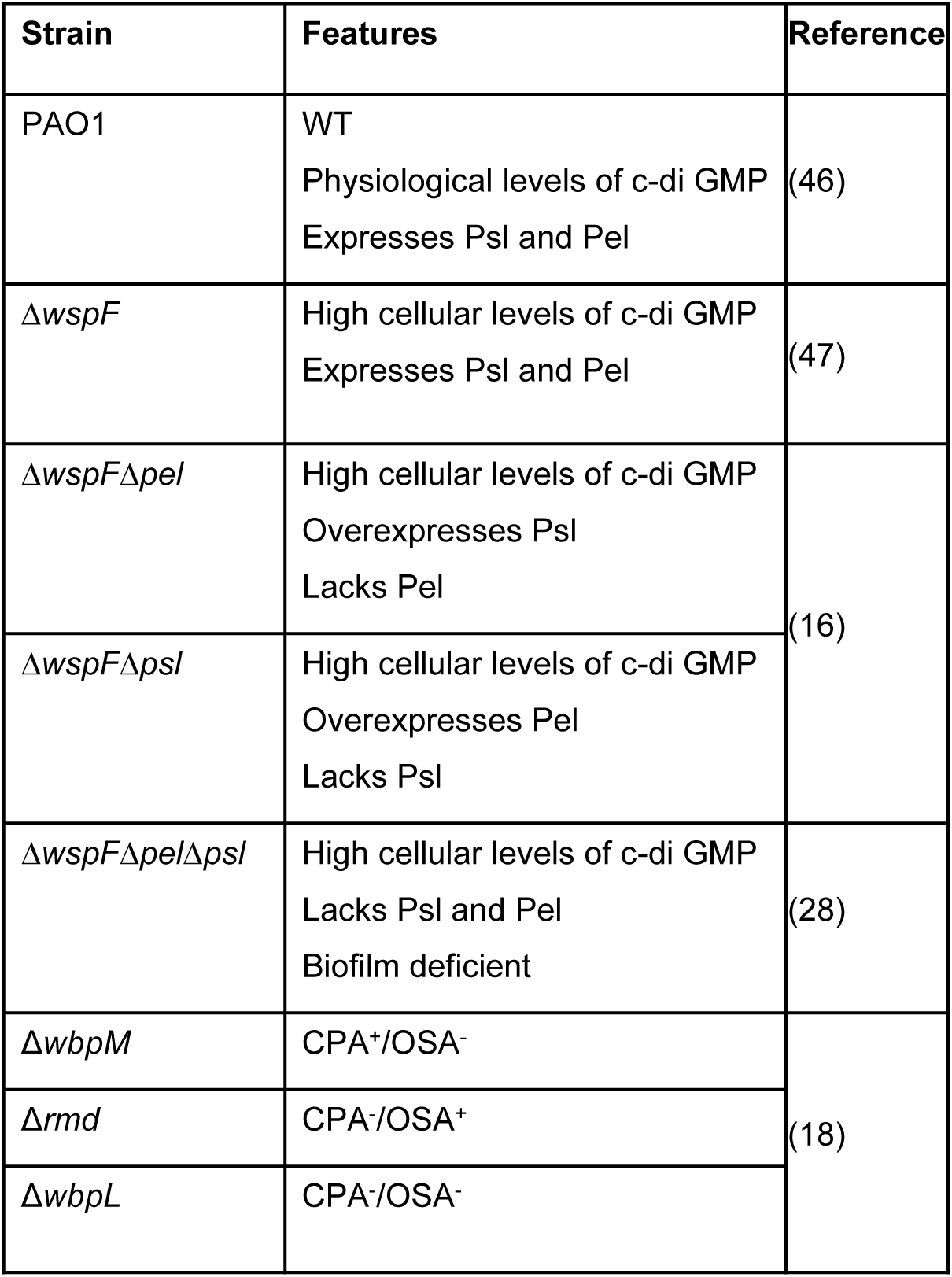
Strains used in this study.

**Figure 3.**
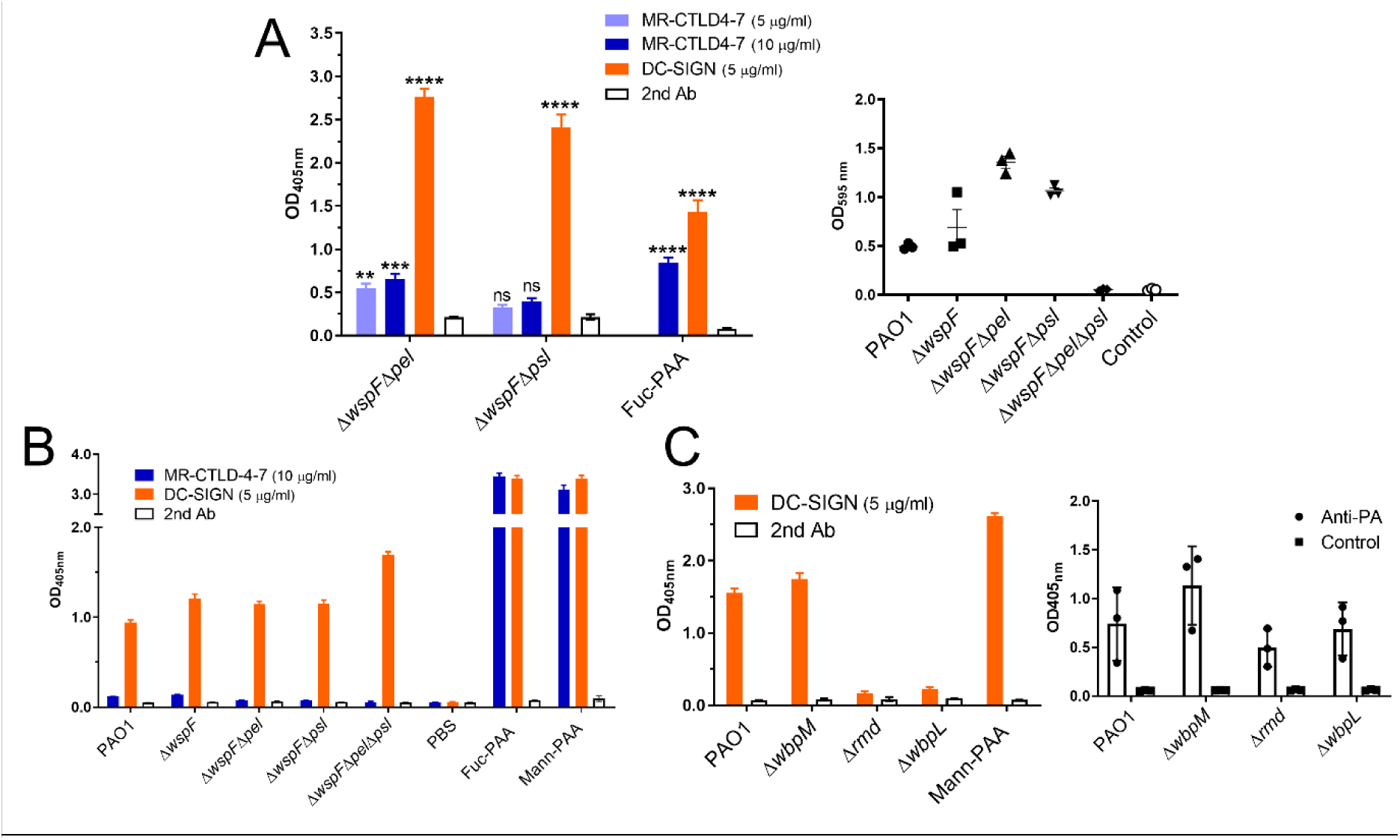
DC-SIGN binds to *P. aeruginosa* biofilms and planktonic cells. **A.** Binding of MR-CTLD4-7 but not DC-SIGN to PAO1 biofilms depends on the presence of the *psl* operon. Biofilms formed by Δ*wspF* Δ*pel* (Psl+/Pel-) or Δ*wspF* Δ*psl* (Psl-/Pel+) were generated in 96 well plates for 24 h, fixed and incubated with MR-CTLD4-7-Fc or DC-SIGN-Fc followed by anti-human Fc antibody conjugated to alkaline phosphatase. Two-way ANOVA with Dunnett’s multiple comparison test. N=3 in triplicate. Right panel. Biofilms formation was confirmed using crystal violet assay. **B.** Planktonic cultures of *P. aeruginosa* PAO1 and different mutants were collected, fixed and used to coat wells of MaxiSorp plates. Wells were incubated with MR-CTLD4-7-Fc or DC-SIGN-Fc followed by anti-human Fc antibody conjugated to alkaline phosphatase. DC-SIGN, but not MR, bind planktonic bacteria and binding is independent from the presence of Psl and/or Pel. N=4 in triplicate. **C.** DC-SIGN binding to planktonic bacteria is dependent on presence of CPA LPS which is absent in the Δ*rmd* and Δ*wbpL* mutants. N=3 in triplicate. Man-PAA and Fuc-PAA refer to commercial mannose and fucose polymers. Right panel: Adherence of planktonic cells to the wells was confirmed by ELISA using an antibody against *P. aeruginosa* (Anti-PA). N=3 in triplicate. Graphs show mean +/− SEM.

### MR and DC-SIGN bind to carbohydrates produced by *P. aeruginosa* biofilms

To determine whether mannose-rich sugars from *P. aeruginosa* biofilms bound MR and DC-SIGN, carbohydrates from cultures of the Pel-deficient mutant Δ*wspF* Δ*pel* (Table 1) were purified as described (19). Two preparations generated independently, 1 and 2, were divided into high (>45 kDa, HMW) and low molecular weight (<45 kDa, LMW) by gel filtration chromatography based on protein standards (19). Gel permeation chromatography (GPC) confirmed differences in size (15,370 Da for LMW-1 and 182,300 Da and 132,670 Da, for HMW1 and HMW-2, respectively. LMW-2 was not investigated) (Figure 4A). A substantial amount of the material in all the samples (~33 – 40% of the total mass) eluted with the included volume. In our system, this means compounds with low MW, i.e. 1000 Da. Their nature is unknown, but we propose that they could be carbohydrate breakdown products. No major protein or DNA contamination were detected based on Silver (Figure S3) and Coomassie staining, protein quantification and spectrophotometry (data not shown). DC-SIGN and MR-CTLD4-7 bind to HMW and LMW preparations (Figure 4B and C) but binding to HMW was stronger. In contrast to their biofilm binding ability, MR-CTD4-7 and DC-SIGN bind similarly to both HMW preparations.

**Figure 4.**
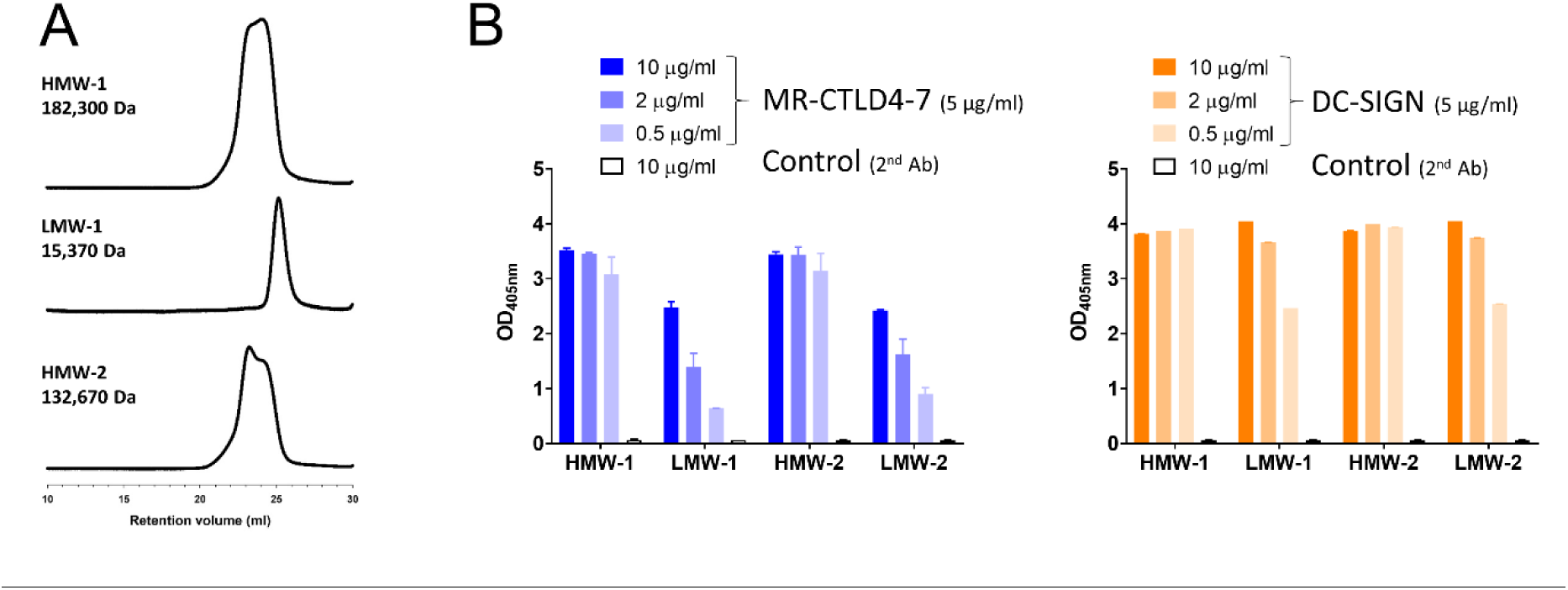
Size analysis and binding to DC-SIGN and MR-CTLD4-7 of *P. aeruginosa* biofilm-associated carbohydrate. **A.** GPC analysis of HMW-1, LMW-1 and HMW-2 confirms successful fractionation into high and low MW forms. HMW-2 contained two peaks poorly resolved. This indicates that this sample is comprised of two components that are similar in MW and perhaps conformation. The MW for HMW-2 reflects the average MW for the entire sample. **B.** Lectin binding assays demonstrate binding of MR-CTLD4-7 and DC-SIGN to HMW-1, LMW-1, HMW-2 and LMW-2. Robust binding of HMW-1 and 2 was observed at 0.5 µg/ml while binding of LMW-1 and 2 at this concentration was substantially reduced. Dose-dependent binding of HMW-1 and HMW-2 to MR-CTLD4-7 and DC-SIGN occurs at lower doses (Figure S4). Fc-chimeric proteins and anti-human Fc-secondary antibody conjugated to alkaline phosphatase were used. Graphs show mean ± SEM of 2 independent repeats done in duplicate.

Initial ^1^H-NMR analysis indicated increased level of impurities in LMW-1 compared to HMW-1 and HMW-2 (data not shown), hence further work largely focused on HMW preparations. The hydrolysed carbohydrate monomer compositions in weight % for HMW-1 is 74.9% mannose, 14.7% glucose, 7.4% galactose, and 3.0% rhamnose and for HMW-2 80.9% mannose, 11.0% glucose, 2.3% galactose, and 5.7% rhamnose. The ^1^H-NMR spectra of HMW-1 and HMW-2 are very similar (Figure 5A) and show that, mannose, the major monomer present, arose from mannan segments in the polymer (Figure S5) (20). The mannose-rich composition of HMW preparations agrees with previous findings (9) and is supported by its recognition by Hippeastrum Hybrid Amaryllis (HHA) lectin (Figure 5B) commonly used to detect the mannose-rich carbohydrate Psl within *P. aeruginosa* biofilms (21). Binding of DC-SIGN and MR to HMW-2 was further confirmed using bio-layer interferometry and purified full-length human MR and biotinylated tetrameric DC-SIGN (22). Analysis of the binding kinetics revealed that both receptors bound HMW-2 with K_D_s in the nM range (Figure 5C).

**Figure 5.**
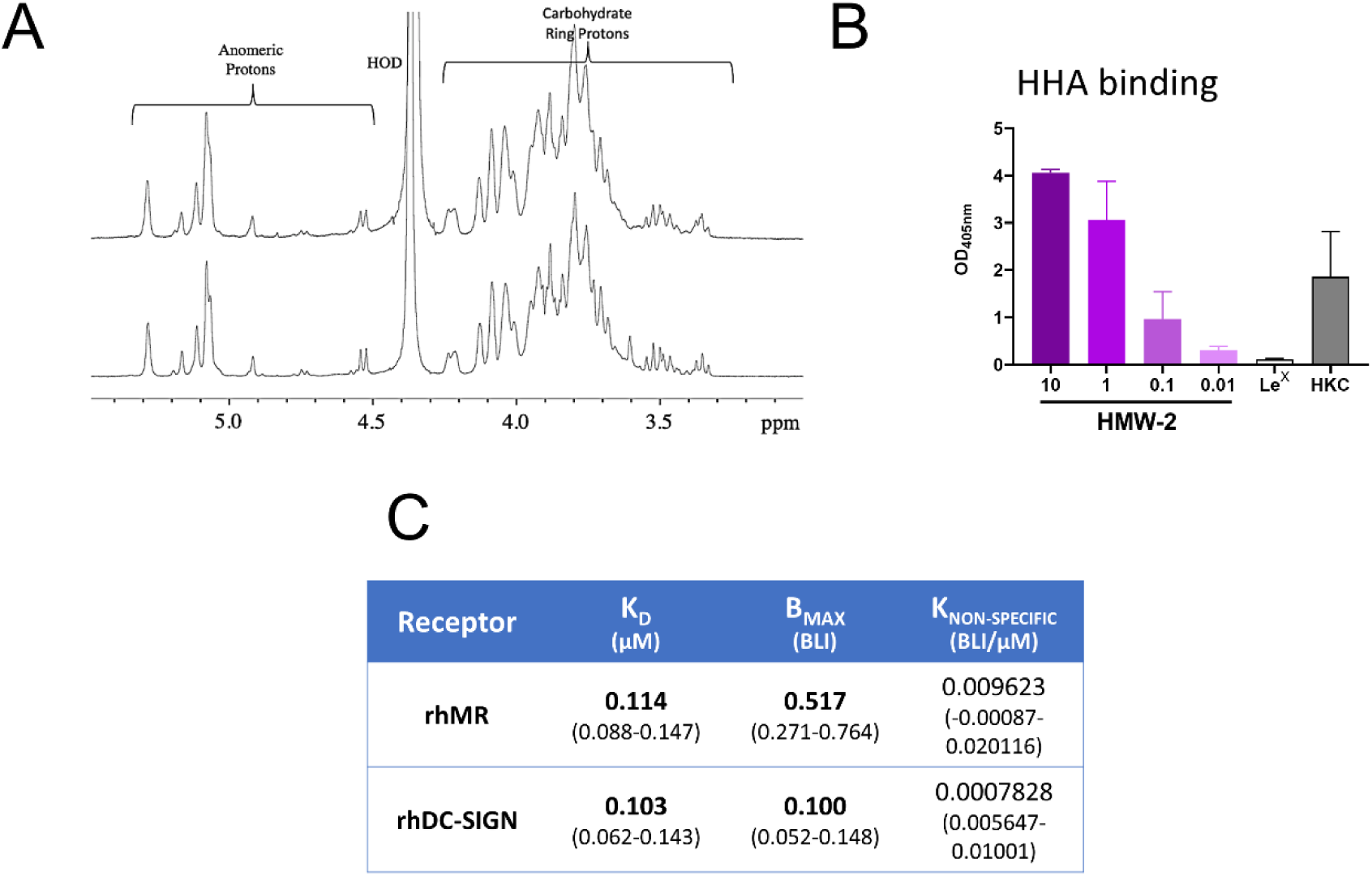
Characterisation of high molecular weight biofilm carbohydrates. **A.** ^1^H-NMR spectra from HMW-1 (top) and HMW-2 (bottom) demonstrate that they are very similar and contain primarily carbohydrates composed of α(1-6) linked mannose segments. **B.** HMW-2 is recognised by the mannose-specific lectin HHA in a lectin binding assay. HHA recognises both (1-3) and (1-6) α-linked mannose structures. LPS-free HMW-2 (Figure S3) and HHA conjugated to alkaline phosphatase were used in these assays. Lewis^x^-PAA and Heat-Killed *Candida albicans* (HKC) were negative and positive controls, respectively. Graph shows Mean ± SEM of two independent repeats done in duplicate. **C**. HMW-2 binds rhDC-SIGN and rhMR. Tetrameric hDC-SIGN, biotinylated and immobilised on a streptavidin sensor and rhMR immobilised on a Ni-sensor were incubated with different HMW-2 concentrations. The table shows equilibrium dissociation constants for the receptor ligand interaction in μM (K_D_); receptor density on the biosensor surface (BMAX) and non-specific binding (K_NON-SPECIFIC_). 95% Confidence intervals in µM are shown within brackets.

### Effect of biofilm carbohydrate on human dendritic cells

Following on previous findings, we next explored the possibility of the mannose-rich HMW biofilm carbohydrate preparations altering the phenotype of human moDCs (MR^+^, DC-SIGN^+^ cells, Figure S6). However, silver staining of HMW-1 and HMW-2 highlighted substantial endotoxin contamination (Figure S3A). Accordingly, HMW-1 and HMW-2 induced high levels of TNF-α by moDCs that were reduced in the presence of polymyxin B. In addition, the pattern of cytokines produced by moDCs in response to HMW-2 was indistinguishable from that of purified endotoxin based on a cytokine microarray assay (Data not shown). LPS removal from HMW-2 was achieved using an endotoxin removal column and confirmed using SDS-PAGE (Figure S3B) and toll-like receptor 4-reporter cells (Data not shown). LPS-free HMW-1 and HMW-2 both retained the ability to bind DC-SIGN and MR-CTLD4-7 (Figure S3C). LPS-free HMW-2 (10, 1, 0.1 µg/ml) in isolation did not induce cytokine production by moDCs nor modified the response of moDCs to purified *E. coli* LPS (Figures S7 and S8). Similarly, LPS-free HMW-2 did not affect the cytokine response of moDCs to Δ*wspF* Δ*psl* biofilms or Δ*wspF* Δ*psl* Δ*pel* cultures (Figure S9). To establish whether HMW-2 modulated other aspects of moDCs biology, we investigated morphological changes in moDCs and DC-SIGN surface distribution after incubation on HMW-2-coated surfaces. Both HMW-2 and LPS-free HMW-2 were tested. moDCs cultured on LPS-free HMW-2 for 24 h display a rounder morphology, characterised by an increased circularity index and reduced perimeter (Figure 6), suggesting changes in the cytoskeleton related to maturation state. These morphological changes were less apparent when using crude HMW-2 indicating that LPS can partially reverse this effect. Analysis of DC-SIGN surface expression showed reduced DC-SIGN labelling (Raw Integrated Density) in cells cultured in the presence of HMW-2 (both crude and LPS-free preparations). When adjusting for cell perimeter (Signal per Unit Area), only the LPS-containing samples showed reduced surface DC-SIGN labelling which is compatible with a more classical moDC activation (23). These findings support the ability of biofilm-associated carbohydrates to influence moDC function in the absence of LPS.

**Figure 6.**
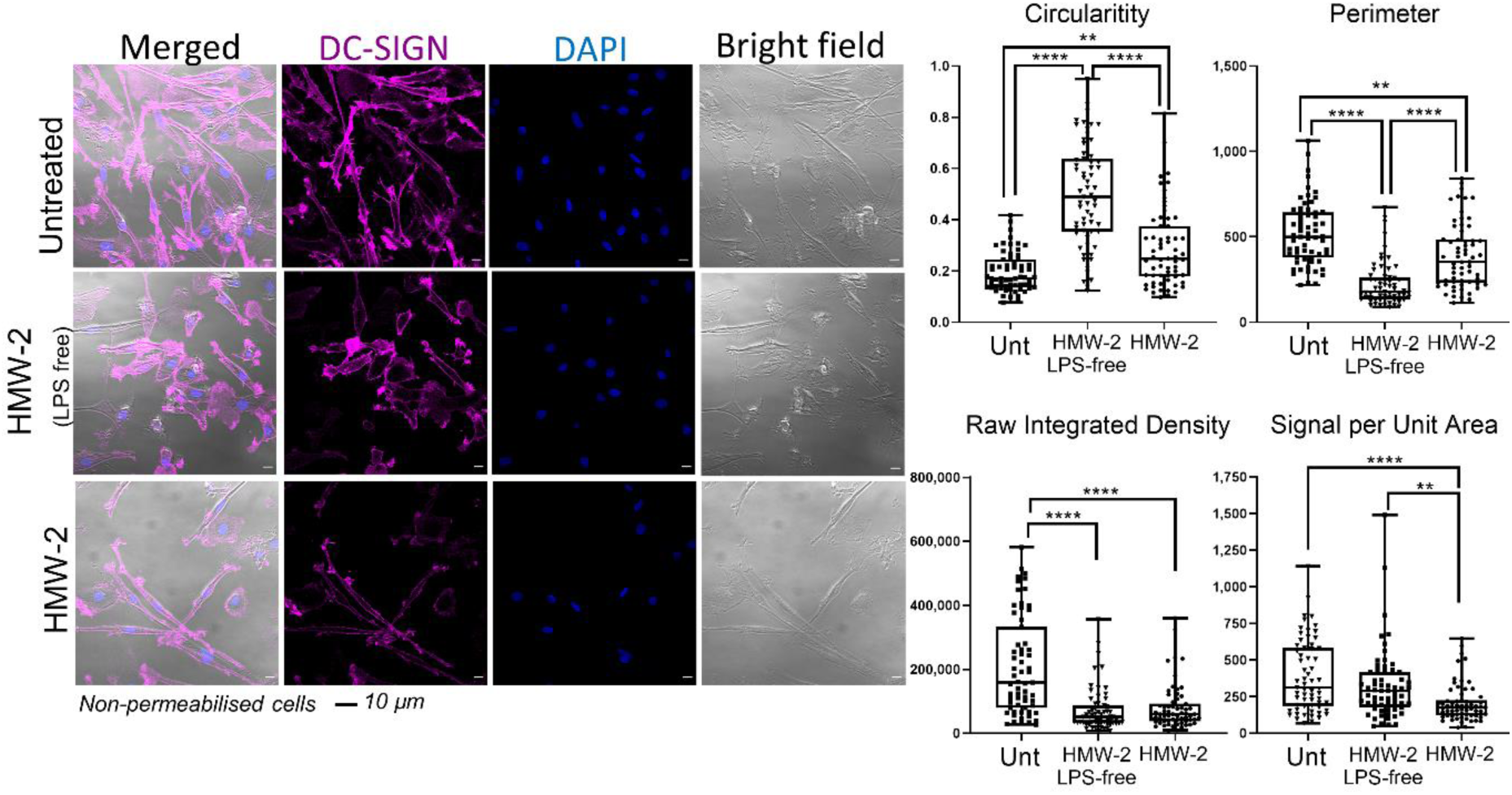
Changes in human dendritic cell morphology in the presence of biofilm-associated carbohydrate. HMW-2 (with and without LPS contamination) diluted in X-vivo-15 (10 µg/ml) were used to coat chambers of µ-slide VI 0.4 flow slides over night at 4°C. moDCs were added (5 × 10^5^ cells per channel) and incubated for 24 h. Samples were then fixed and stained for DC-SIGN (magenta) and nucleus (DAPI, blue). The figure shows representative images from unpermeabilised samples. Permeabilised samples, including secondary antibody control are shown in Figure S10. Cells were analysed for changes in shape (Circularity Index), size (Perimeter) and DC-SIGN labelling intensity (Raw Integrated Density and Signal per Unit Area). Data derive from 3 independent experiments, 20 cells per experiment were analysed. Statistical significance assessed using Kruskal-Wallis test corrected for multiple comparison using a Dunn’s multiple comparison test.

### Mannose-rich biofilm carbohydrate interferes with the function of cell-associated DC-SIGN and MR

MR and DC-SIGN are important endocytic receptors expressed by antigen presenting cells. Therefore, in a different set of experiments we tested whether HMW-1 and HMW-2 could interfere with their endocytic activities. moDCs internalise fucose-PAA-FITC (model ligand for MR and DC-SIGN), Lewis^x^-PAA-FITC (model ligand for DC-SIGN) and, poorly, galactose-PAA-FITC (not recognised by MR or DC-SIGN). Presence of HMW-1 and HMW-2 (crude preparations) partially inhibits uptake of Lewis^x^-PAA-FITC but not that of fucose-PAA-FITC or galactose-PAA-FITC by moDCs (Figure 7A). These findings were not affected by Polymyxin B (100 µg/ml) (Figure S11) suggesting that these observations are LPS-independent. These results indicate specific ability of HMW-1 and HMW-2 to modulate DC-SIGN-mediated endocytosis in moDCs. We next employed cell lines expressing either DC-SIGN (U937-DC-SIGN) or MR (CHO-MR) to validate these findings. U937-DC-SIGN cells associate with Lewis^x^-PAA-FITC and fucose-PAA-FITC specifically and HMW-1 and HMW-2 inhibit both activities (Figure 7B) indicating that biofilm carbohydrates can interact with cell-associated DC-SIGN and compete with DC-SIGN ligands for binding. CHO-MR internalise fucose-PAA-FITC and, weakly, Lewis^x^-PAA-FITC (Figure 7C). Uptake of Lewis^x^-PAA by MR was unexpected as the MR-CTLD4-7 fragment does not bind Lewis^X^ in ELISA-based assays (Data not shown) and this sugar lacks the sulphated moiety required for binding to MR-CR domain (24). Surface plasmon resonance (SPR) analysis using full length human MR confirmed the capacity of MR to bind Lewis^x^ polymers (Figure S12) supporting that uptake of Lewis^X^ by CHO-MR cells is MR-mediated. HMW-1 and HMW-2 interfere with uptake of Lewis^X^-PAA-FITC, but not fucose-PAA-FITC, by CHO-MR cells. LPS removal does not affect these findings (Figure S13). These results suggest that HMW biofilm carbohydrates interfere with uptake of selected MR ligands, but possibly only those with lower binding avidity. No sugar uptake was observed in the case of U937 or CHO control cells (Data not shown). Combined these data suggest that HMW biofilm carbohydrates interfere with MR and DC-SIGN function and unveil the differential contributions of MR and DC-SIGN to sugar uptake in human moDCs with fucose being preferentially internalised through MR (not inhibited by HMW biofilm carbohydrates) and Lewis^X^ by DC-SIGN and/or MR (both inhibited by HMW biofilm carbohydrates).

**Figure 7.**
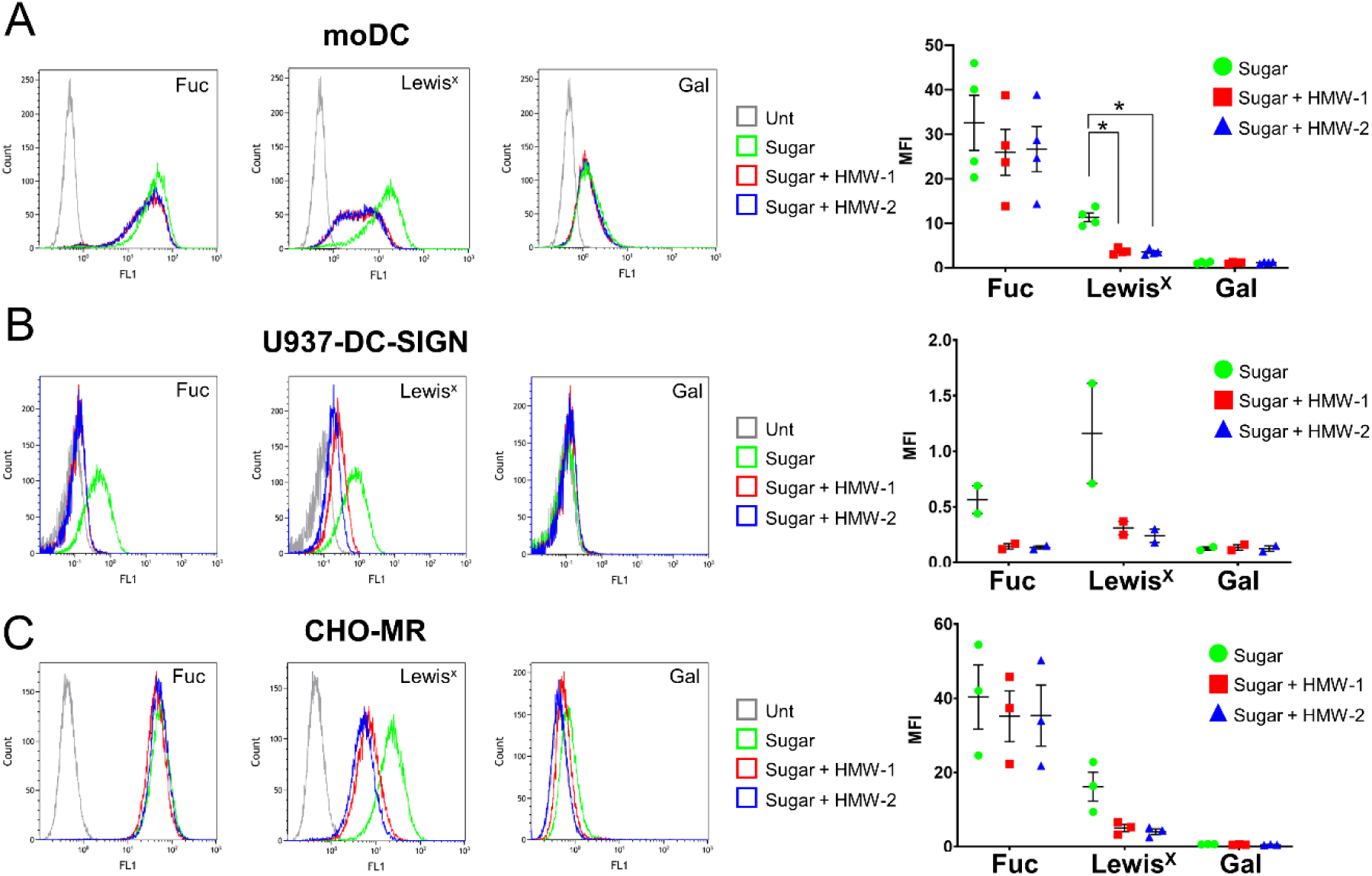
Inhibition of sugar uptake by MR and DC-SIGN expressing cells by HMW-1 and HMW-2. Cells were treated with HMW-1 or HMW-2 (10 µg/ml) for 1 h, then fluorescently labelled sugars were added for a further 1 h. Association of fluorescent sugars to cells was measured by flow cytometry. Representative histograms and scatter plots depicting mean ± SEM of median fluorescence intensity (MFI) are shown for each cell type. **A.** Human moDCs (expressing MR and DC-SIGN) internalise fucose (Fuc), Lewis^x^ and galactose (Gal) PAA-FITC polymers but only Lewis^x^ uptake is reduced by HMW-1 and HMW-2, N=4. **B.** U937-DC-SIGN cell line (express DC-SIGN, but not MR) internalise fucose and Lewis^x^ polymers and their uptake is reduced by HMW-1 and HMW-2, N=2. **C.** CHO-MR cells (express MR, but not DC-SIGN) internalise fucose and Lewis^x^ polymers and only Lewis^x^ uptake is reduced by HMW-1 and HMW-2, N=3. Repeated-measures ANOVA was performed to determine statistical significance; *, ≤0.05. Fuc: fucose; Gal: galactose.

## Discussion

In this study, we demonstrate (i) direct recognition of *P. aeruginosa* biofilms by the C-type lectin receptors DC-SIGN (CD209) and MR (CD206) and detect CPA-LPS-dependent binding of DC-SIGN to planktonic PAO1 cells; (ii) describe the composition and structure of HMW carbohydrate preparations from *P. aeruginosa* biofilms; (iii) show direct binding of DC-SIGN and MR to biofilm-associated carbohydrates; and (iv) provide evidence for changes in human DC morphology and DC-SIGN and MR endocytic activity in the presence of biofilm-associated carbohydrates. The key message of these studies is that, at least under our experimental conditions, biofilms display carbohydrate ligands for immune C-type lectin receptors that could contribute to the modulation of immunity towards these structures.

Biofilms are major drivers of bacterial pathogenesis in the context of chronic infections. Historically, protection against immune attack alongside antibiotic tolerance, were postulated as the key advantages conferred by these biofilms during infection but research into their role in modulating immunity is gathering momentum (5, 25). In the context of infection, MR and DC-SIGN are considered promoters of regulatory immune mechanisms designed to curtail damaging inflammatory processes and MR and/or DC-SIGN binding are viewed as immune evasion mechanism(s) (11, 12, 26, 27). Neither DC-SIGN or MR display canonical signalling motifs at their cytosolic domains and instead of triggering cellular responses modulate the outcomes to stimulation of signalling pattern recognition receptors such as Toll-like receptors (11, 12, 27). Carbohydrates are molecular patterns not normally associated with bacterial infections but their abundance in bacterial biofilms necessarily alters this perception. Fungal pathogens and viruses, together with *Mycobacterium tuberculosis*, display ligands for C-type lectins and some of their sugar-bearing structures, such as mannan, β-glucan, mannose patches and lipoarabinomannan, modulate immunity through lectin engagement. Since biofilms are largely associated with chronic disease, it is plausible to speculate that after initial infection mediated by planktonic-like cells, a biofilm-like lifestyle could both counterbalance immune attack while gearing immunity towards non-resolving, ineffective immunity (Figure 8).

**Figure 8.**
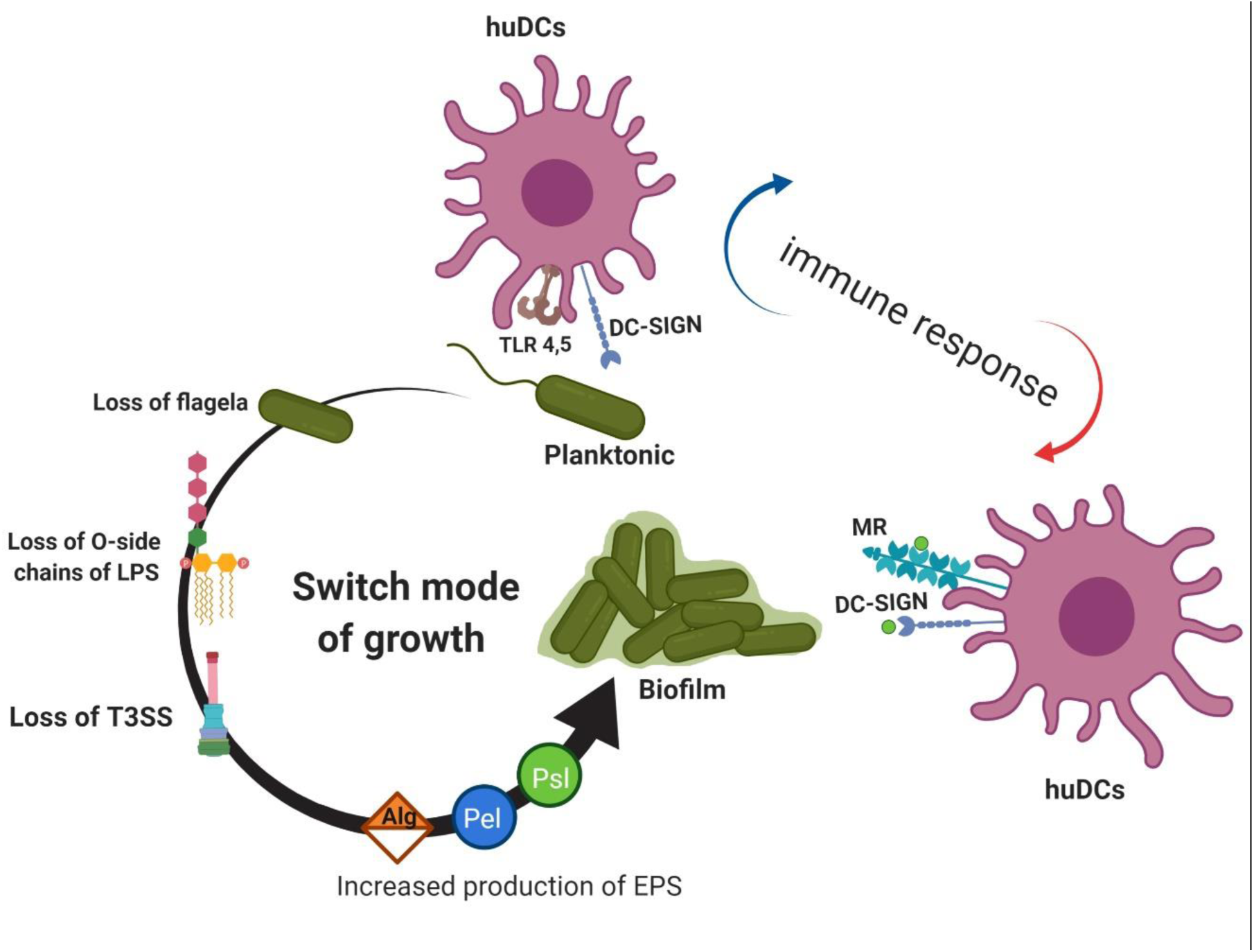
Role of lectin receptors during *P. aeruginosa* infection. Planktonic *P. aeruginosa* and biofilms co-exist in the host, with planktonic bacteria primarily associated with acute infections and biofilms with chronic infections. Planktonic bacteria display traits associated with enhanced cytotoxicity (T3SS+) and ability to stimulate immune cells (Flagellin+) and can trigger multiple signalling pathways through engagement of pattern recognition receptors (Toll-like receptors are displayed as example). In this instance DC-SIGN, could modulate cellular activation and lead to upregulation of IL-10 production (as observed in other pathogens). Biofilms engage both DC-SIGN (through biofilm-associated carbohydrate and ligands in planktonic cells) and MR (through biofilm-associated carbohydrates). In addition, biofilm-associated bacterial cells could display reduced ability to cause cytotoxicity and stimulate pattern recognition receptors. In this instance MR and DC-SIGN could modulate an already altered cellular activation likely leading to further modulation of immunity away from pro-inflammatory Th1-dominated responses.

Both HMW preparations investigated in this study are mannose-rich and contain α-mannose segments. Despite the clear differences observed in DC-SIGN and MR binding to whole biofilms, their comparable binding to purified HMW preparations indicates that the purification procedure likely selects for common MR and DC-SIGN ligands within the biofilm structure. In addition there is a possibility for soluble bacterial proteins such as CrdA, shown to reinforce biofilm structure through Psl binding, to block the MR and/or DC-SIGN binding sites within the biofilm structure (28). Finally our results agree with work implicating MR as receptor for Slime-GLP, a crude ethanol extract of *P. aeruginosa* biofilm matrix (29).

Binding of DC-SIGN and MR to *P. aeruginosa* biofilms was detected using ELISA-based assays and confocal microscopy and there was good correlation between both assays with DC-SIGN binding being more abundant and widely distributed than that of MR. Findings agree with the broader binding specificity observed for DC-SIGN (recently reassessed in (30)). We hypothesise that MR binds a subset of the DC-SIGN binding sites as suggested by the clustering of the binding sites for both lectins. The confocal study also supports the heterogeneity of mannose-rich structures within *P. aeruginosa* biofilms as both MR and DC-SIGN binding patterns differ from that of the HHA lectin, normally used for the detection of mannose-rich biofilm-associated carbohydrates (21) (Compare Figure 2 and Figure S2). HHA preferentially binds bacterial aggregates, which indicates preferential binding to mannose structures associated to bacterial cells. It is possible that MR and/or DC-SIGN binding sites in cell-associated carbohydrates are blocked through binding to the mannose-specific *P. aeruginosa* lectin LecB that directly interacts with Psl (31). Our results agree with the existence of distinct Psl epitopes (class I, II, and III) which can be targeted with different monoclonal antibodies (mAbs) (32) and are differentially distributed within mature PAO1 biofilms (33).

The predicted carbohydrate structure for the HMW preparations used in this work do not conform to that described for Psl, the mannose-rich neutral polysaccharide produced by PAO1 via the *psl* operon products. Byrd et al. described Psl as repeating pentameric units of D-mannose, L-rhamnose and D-glucose (9). In contrast, our preparation contains a small proportion of galactose and, unlike the structure proposed for Psl, lacks mannose β anomers. There is a high proportion of 1-6-linked-α-mannose with some 1-2 linkages, characteristics of mannans. In *C. albicans* the structure of mannan varies depending on the culture conditions (34) and it is highly feasible that differences in growth conditions, purification procedures, including selection of HMW forms, and bacteria strain (WT vs Δ*wspF* Δ*pel*) could account for these observations. In agreement with our findings Bates et al using the same purification procedure as ours (19) identified galactose alongside mannose, glucose and rhamnose in carbohydrates generated from two *P. aeruginosa isolates* (700829 and 700888) and there was abundance of 2-6 linked (32-28%), 2-linked (20-19 %), 3-linked (16%) and terminal (23-27 %) mannose. Hence, this work opens the exciting possibility of mannose-rich carbohydrates in *P. aeruginosa* not conforming to a unique structure but displaying adaptability to environmental changes and/or bacterial genetic makeup further broadening the range of biofilm arrangements and associated immune responses. The strain Δ*wspF* Δ*pel* used to generate the carbohydrates in this study produces high cyclic-di-GMP levels that could impact on the regulation of carbohydrate structures. For instance, McCarthy et al demonstrated regulation of LPS modifications by cyclic-di-GMP in *P. aeruginosa* through binding to WarA, a methyltransferase that regulates O-antigen modal distribution and interacts with components of the LPS synthesis pathway (35).

Release of mannose-rich, well-defined HMW polymeric entities by biofilms (our HMW preparations derive from cell-free supernatants) raises the possibility of biofilm-associated carbohydrates acting as immunomodulators on their own. Assays to date using moDCs were restricted to early responses and failed to detect major changes in cytokine responses to LPS or Psl-deficient biofilms in the presence of LPS-free HMW-2. Future work will focus on functional studies such as ability of moDCs to activate T cells in the presence of HMW. Indeed, the phenotypical changes observed in moDCs with only 10 µg/ml of HMW (cell rounding and reduced Lewis^x^ binding) indicates that HMW could interfere with DC-T cell interactions or DC migration by interfering with ICAM-3 or ICAM-2 adhesion through DC-SIGN (36, 37).

Assays in which moDC where incubated with biofilms with different carbohydrate compositions (PAO-1, Δ*wspF*, Δ*wspF* Δ*pel* and Δ*wspF* Δ*psl*), Figure S14), failed to show selective production of cytokines in response to particular biofilm types. These results indicate that under these experimental conditions the presence and absence of Psl or Pel do not have a major impact on early cytokine production by moDCs. In addition, preliminary results did not to show clear trends when moDC-Δ*wspF* biofilm co-cultures were performed in the presence and absence of blocking antibodies against MR and DC-SIGN. Differences in biofilm formation among assays together with donor variability likely contribute to these findings. While our moDCs consistently expressed MR and DC-SIGN, levels of surface MR in particular, were highly variable among donors (Figure S6). In addition, it is possible that during the fixation process, although bacteria remained damaged and non-culturable for a least 4 h, total bacterial cell death was not achieved, which could promote inflammatory activation. Current work in the laboratory focuses on further optimisation and validation of biofilm-moDC co-cultures and potential stratification of donors based on moDCs receptor expression.

The dominance of DC-SIGN binding to *P. aeruginosa* biofilms and planktonic cells highlights the importance of using infection models where DC-SIGN is present. Early observations linked DC-SIGN expression in DCs, with biofilm positivity in chronic rhinosinusitis with nasal polyposis (38), and suggest unique immune responses in the presence of biofilms that correlate with DC-SIGN expression. In human skin, dermal macrophages express MR and DC-SIGN (39) and both receptors could contribute to immune responses to *P. aeruginosa* wound infections. Similarly alveolar macrophages from people with cystic fibrosis have increased levels of MR (40) and DC-SIGN expression in these cells has also been described in tuberculosis patients (41). Suitable models to establish contribution of DC-SIGN during *P. aeruginosa* infection need to consider lack of DC-SIGN orthologs in mice (12), hence DC-SIGN transgenic mice offer a suitable alternative (42, 43).

In summary, this work demonstrates direct interaction between biofilm-associated carbohydrates and immune C-type lectins and opens the possibility for these receptors to contribute to the establishment of chronic infections.

## Materials and Methods

### Biofilm quantification assay

All strains (Table 1), unless otherwise stated, were grown on Lysogenic Broth (LB) agar plates from glycerol stocks stored at −80°C and incubated overnight at 37°C. Overnight cultures in X-vivo-15 media (Lonza) (5ml, 37°C, 200/220 rpm) diluted to OD_600nm_ 0.01 were cultured for 3 h at 37°C, 200/220 rpm. The OD_600nm_ of mid log phase cultures in X-vivo-15 was adjusted to 0.04 OD_600nm_ and 100 μl of cultures were added into each well of a UV-sterilised 96-well plate [Costar (9017, Corning) or Maxisorp (439454, Nunc immune-plate)]. Cultures incubated for 24 h at 37°C, 5%CO_2_ were washed three times with 200 μl of HPLC water and stained with 125 μl of 1% (w/v) crystal violet (1 h, room temperature (RT)). After washing three times in water, the stain was solubilised by adding 200 μl of 70% ethanol for 15 min; 125 µl was transferred into a clean 96 well Costar plate to measure the absorbance at 595 nm using a Multiskan FC (Thermo Scientific).

### Analysis of the adhesion of planktonic *P. aeruginosa* to plastic

Overnight *P. aeruginosa* cultures were centrifuged at 16,000 × g for 5 min at 4°C, washed twice with PBS, and re-suspended in 4% (v/v) paraformaldehyde (15710-S, Electronic Microscopy Sciences, USA) in PBS for 30 min at 4°C. After fixation, cultures were washed once with PBS, adjusted to 0.5 OD_600nm_ in PBS, and pipetted onto Maxisorp plates (50 µl/well). After washing three times with PBS blocking was carried out by adding 50 µl of 3% (w/v) bovine serum albumin (BSA) (80400-100, Alpha diagnostics) prepared in PBS. Rabbit anti-*P. aeruginosa* polyclonal antibody (50 µl/well, ab68538, Abcam) diluted 1:1000 in PBS was added and incubated for 90 min at RT. After three washes in PBS, the plate was incubated with 50 µl of goat anti-rabbit IgG conjugated to alkaline phosphatase diluted 1:2000 (A3687, Sigma) in PBS for 1 h at RT. After three washes with AP buffer (100 mM Tris-HCl, 100 mM NaCl, 1 mM MgCl2, pH 9.5), 50 µl of p-nitrophenyl phosphate substrate solution (Sigma) were added to each well and incubated for 30-40 min at room temperature in the dark. Absorbance was measured at 405 nm using a Multiskan FC (Thermo Scientific).

### Lectin binding assays

Assays for the binding of chimeric proteins to fixed *P. aeruginosa* biofilms, fixed planktonic *P. aeruginosa* cells and purified carbohydrate were performed as follows. Biofilms were grown on a Costar (9017, Corning) or Maxisorp (439454, Nunc immune-plate) plate over 24 h and fixed with 50 µl of 2% paraformaldehyde in PBS for 10 min at 4°C. For fixed *P. aeruginosa* cells, wells of Maxisorp plates were coated with fixed bacteria (100 µl/per well) and incubated at 4°C overnight. Purified biofilm carbohydrate was added to Maxisorp plates overnight (50 µl/well in 154 mM NaCl, 37°C). In all instances, plates were washed three times with TBS (10 mM Tris-HCl, pH 7.5, 10 mM CaCl_2_, 154 mM NaCl and 0.05% (v/v) Tween 20). Chimeric proteins MR-CTLD4-7 (CTLD-4-7-Fc, prepared in house, (14)) and DC-SIGN (Fc-DC-SIGN-Fc, R&D) (50 µl/well in TBS) were added and incubated for 2 h at RT. After three washes with TBS, anti-human Fc-conjugated to alkaline phosphatase (A9544, Sigma) was added (50 µl/well) and incubated for 1 h, RT (1:1000 dilution). After washing three times with TBS, alkaline phosphatase activity was measured as above. Inhibition assays were carried out as above but using TSB buffer containing 1M NaCl (TSB-high salt). MR-CTLD4-7 and DC-SIGN were pre-incubated with different concentrations of the monosaccharides mannose (63579, Fluka), fucose (47870, Fluka), or galactose (4829, Fluka) in TSB-high salt for 30 min at RT. After pre-incubation, proteins were added to appropriate wells containing biofilms. Polymers containing D-mannose, L-fucose, Lewis^x^ or D-galactose (2-5 µg/ml, 50 µl per well, Lectinity) were used as controls. Binding of HHA to purified carbohydrates was tested in a similar way using alkaline phosphatase-conjugated HHA (20 µg/ml, LA-8008-1 EY laboratories).

### Study of DC-SIGN and MR-CTLD4-7 binding to *P. aeruginosa* biofilms by confocal microscopy

Biofilms generated under flow on µ-Slide VI 0.4 (Ibidi) as described in Figure S2, were fixed with 4% (v/v) paraformaldehyde in PBS (100 µl per channel, 10 min, 4°C), and washed three times with TSB buffer. In some instances, wells were stained with FM 1-43 FX membrane dye (100 µl per channel, 2-10 µg/ml, F35355, Thermofisher) in PBS for 30 min on ice. Following three washes with TSB buffer MR-CR-FNII-CTLD1-3 (15), MR-CTLD4-7 or DC-SIGN (30 µl per channel, 10 µg/ml in TSB buffer) were added and incubated for 2 h at RT. After washing three times with TSB buffer, 100 µl of TSB containing 10 µg/ml goat anti-Human IgG conjugated to Alexa fluor 647 (A21445, Invitrogen) and 3%(v/v) Donkey serum (D9663, Sigma) in TBS were added and incubated for 1 h at RT. Following three washes with TSB, DNA was labelled with DAPI (100 µl per channel, 2 µg/ml D9542, Sigma-Aldrich) in PBS for 15 min, RT. The plates were washed with TSB and mounted in Ibidi mounting media (50001, Ibidi, 50001) before storing at 4°C in the dark. Confocal images were acquired using Zeiss LSM 880 using a 40x /1.20 water objective, the collection was not done with filters. Fluorescence emission was collected between 434 and 515 nm (DAPI), 469-538 nm (FM 1-43FX), 641-688 nm (AF 647). Stack size (49.43 µm, y: 49.43 µm, z: 7.5-12.9 µm). Presented single slice size (49.43 µm, y: 49.43 µm, z: 0.288-0.293 µm). Images were processed using Fiji (44).

### Carbohydrate purification

Carbohydrate was extracted from ∆*wspF* ∆*pel* (7) cultures as described previously (19). Overnight cultures in 20 ml, TSB medium, (22092, Sigma) were added to TSB medium in a 1.5 L flask (400 ml per flask) and incubated statically at 37°C for five days. Cultures were treated with formaldehyde (final concentration 0.02% (v/v) from 36.5% solution-33220, Sigma-Aldrich) 1 h, RT, 100 rpm followed by NaOH (a final concentration of 275 mM, S318-1, Fisher Scientific) 3h, RT, 100 rpm. Cultures were centrifuged (16,000 x g, 1 h, 4°C) and supernatant was collected, filtered and dialysed/concentrated against HPLC water using VIVAFLOW 200, MWCO 10 kDa (Sartorius Stedim Biotech) to a maximum final volume of 50 ml. Proteins and nucleic acids were precipitated using tri-chloro-acetic acid (TCA, 20% (w/v), 3000-50, Fisher scientific) at 4°C for 30 min. The solution was centrifuged (16,000 x g, 1 h at 4 °C) and the supernatant was collected into a fresh clean glass bottle and EPS was precipitated away from lipids by incubation with 1.5 volumes of cold 95% (v/v) ethanol −20 °C, 24 h. This step was done twice to improve purity. The solution was centrifuged at 16,000 x g for 1 h at 4°C and the pellet was re-suspended in HPLC water, dialysed against HPLC water using a 12–14 kDa MWCO membrane (68100, Snakeskin), and lyophilized. The lyophilized powder was re-suspended in PBS (pH 7.4) and fractionated on a HiPrep 26/60, Sephacryl S-200 HR gel filtration column (GE Healthcare) calibrated with protein standards (1511901, Bio-rad) to generate a standard curve showing the retention time of molecular weight standard. For the gel filtration, the equivalent of 2.4 or 3.6 litres of culture were pooled for each column run. High molecular weight (>45 kDa) and low molecular weight preparations (<45 kDa) were pooled, dialysed against HPLC water. Endotoxin was eliminated by repeated passages (x10) through an endotoxin removal column (Thermofisher UK) following the manufacturer’s recommendations.

### Carbohydrate analysis

#### Molecular weight analysis

Molecular weight, polydispersity and polymer distribution values were derived from gel permeation chromatography (GPC) with a Viscotek/Malvern GPC system consisting of a GPCMax autoinjector fitted to a TDA 305 detector (Viscotek/Malvern, Houston, TX). The TDA contains a refractive index detector, a low angle laser light scattering detector, a right angle laser light scattering detector, an intrinsic viscosity detector and a UV detector (λ = 254 nm). Three Waters Ultrahydrogel columns, i.e. 1200, 500 and 120, were fitted in series (Waters Corp. Milford, MA). The columns and detectors were maintained at 40°C within the TDA 305. The system was calibrated using Malvern pullulan and dextran standards. The mobile phase was 50mM sodium nitrite. The mobile phase was filter sterilized (0.45 µm) into a 5 L mobile phase reservoir. Psl samples were dissolved (6 mg/ml) in mobile phase (50 mM sodium nitrite, pH 7.3). The samples were incubated for ~60 min at 60°C, followed by sterile filtration (0.45 μm) and injected into the GPC (50 - 200 μL). Sample recovery was routinely >90%. Dn/dc for each sample was calculated using the OmniSec software (v. 4.6.1.354). The data were analysed using a single peak assignment in order to obtain an average molecular weight for the entire polymer distribution. Each sample was analyzed in duplicate or triplicate. Replicate analysis of calibration standards indicated reproducibility of + 3%, which is well within the limits of the technique.

#### Proton nuclear magnetic resonance

Carbohydrate preparations were structurally characterised by solution-state 1D 1H NMR spectroscopy and 2D COSY and HSQC NMR spectroscopy. 1D NMR data acquisition and analysis was based on methods from Lowman et al. (20, 34). Briefly, NMR spectra were collected on a Bruker Avance III 400 NMR spectrometer operating at 331oK (58°C) in 5mm NMR tubes. Each carbohydrate (about 5 mg) was dissolved in about 550 µl DH_2_O (Cambridge Isotope Laboratories, 99.8+% deuterated). Chemical shift referencing was accomplished relative to TMSP at 0.0 ppm. The proton 1D NMR spectra were collected with 36 scans, 65,536 data points, 20 ppm sweep width centered at 6.2 ppm, and 1 s pulse delay and processed using exponential apodization with 0.3 Hz line broadening. COSY spectra were collected using 2048 by 128 data points, 16 dummy scans, 64 scans, and 9.0 ppm sweep width centered at 4.5 ppm and processed with sine apodization in both dimensions and zero-filled to 1024 data points in f1. HSQC spectra were collected using 1024 by 256 data points, 4 dummy scans, 128 scans, and 6.0 ppm sweep width in f2 and 185 ppm sweep width in f1 and processed with qsine apodization in both dimensions and zero-filled to 1024 data points in both dimensions. Polymer hydrolysis was accomplished by heating the isolate in 33% TFA-d in D2O at 80 °C overnight. Processing was accomplished with Bruker TopSpin (version 4.0.6) on the MacBook Pro.

### Measurement of HMW-2 binding to recombinant human MR and biotinylated DC-SIGN by biolayer interferometry

Binding experiments were performed on an Octet K2 biolayer interferometry system (ForteBio, San Jose, CA) in 10X Kinetics Buffer at 30°C and 1000 RPM. Black bottomed 96-well plates were from Grenier Bio-One and the optical biosensor probes from ForteBio. Recombinant human MR (CD206) with a poly-His-tag was purchased from R and D Systems (Minneapolis, MN). Biotinylated DC-SIGN was generated as described (22). Ni-NTA or SA biosensor was placed in the instrument, and after an equilibration period of 3 min the biosensor was exposed to either the poly-His-tagged MR or the biotinylated DC-SIGN at a concentration of 0.1 mg/mL for 5 min, and then transferred to 10 X Kinetics Buffer for 10 min to establish the BLI signal from the immobilized receptor. Following this, a series of eight similar 5 min exposures to 2-fold increasing concentrations (3.125-400 mg/mL) of carbohydrate each followed by 10 min dissociation in 10X Kinetics Buffer were performed. For each exposure, the equilibrium BLI signal was measured 20 s after the switch to Kinetics Buffer and used in the analysis of binding. A parallel biosensor with immobilized receptor, but not carbohydrate, was placed in 10X Kinetics Buffer and used to control for receptor dissociation during the experiment. Data analysis was performed on GraphPad Prism 7.0 software. The series of BLI signals for each concentration was fit to models of nonspecific linear binding, saturable specific binding, and specific binding plus nonspecific binding with either local variable or shared global values for each parameter. The sequential F-statistic with a p<0.05 was used to select the most appropriate model for each carbohydrate – receptor interaction. Results are reported as mean values with 95% confidence intervals for apparent equilibrium dissociation constant (K_D_), maximum BLI signal (B_MAX_), and nonspecific binding (K_NS_).

### Generation of human monocyte-derived dendritic cells

Human monocyte-derived dendritic cells (moDCs) were prepared from buffy coats (Blood Transfusion Service, Sheffield, UK). Use of buffy coats was approved by the Faculty of Medicine and Health Sciences Research Ethics Committee. PBMCs were isolated using Histopaque-1077 (H8889, Sigma) and monocytes were isolated using human CD14 MicroBeads (130-050-201, Miltenyi Biotec) following the manufacturer’s protocol. Purified monocytes were cultured in RPMI complete medium [RPMI-1640 (R0883, Sigma), 15% (v/v) human AB serum (PAA Laboratories, UK), 2 mM L-glutamine (G7513, Sigma), 10 mM HEPES (15630056, Gibco), 50 ng/ml recombinant human granulocyte macrophage colony–stimulating factor (rhGM-CSF) (130-093-865, Miltenyi Biotec), and 50 ng/ml rh interleukin (IL)-4 (130-093-921, Miltenyi Biotec)] and cultured at 37°C, 5%CO_2_ for 6-7 days. On Day 3, fresh RPMI complete media containing growth factors was added to each well.

### Analysis of changes in moDC morphology after incubation with HMW

For the analysis of morphological changes in moDCs in the presence of Psl, µ-Slide VI 0.4 tissue culture treated channels (80606, Ibidi) were coated with HMW (10 µg/ml, in X-vivo-15) overnight at 4°C. After washing, and addition of moDCs (5 x 104 cells per channel in X-vivo 15 containing GM-CSF and IL-4) cultures were incubated for 24 h at 37°C, 5% CO_2_. Samples were fixed in 4% (v/v) Paraformaldehyde Aqueous Solution (15710-s, EM Grade) in PEM (Cytoskeleton-preserving buffer PIPES-EGTA-Magnesium, 80 mM PIPES pH 6.8, 5 mM EGTA, 2 mM MgCl_2_) for 15 min, RT. Slides were washed in PEM and blocked with 5% donkey serum in PEM in the presence or absence of 0.25% Triton X-100. After 3 washes in PEM, cells were labelled with DAPI (1.5 µg/ml in PBS, D9542, Sigma-Aldrich), washed in PBS and mounted in Ibidi mounting media (50001, Ibidi). Confocal images were acquired using Zeiss LSM 710 under 40x /1.3 oil objective, without filters. Fluorescence emission was collected between 410 and 585 nm (DAPI) and 638-755 nm (AF 647). Image size (212.55 µm, y: 212.55 µm, z: 0.406 µm). Images were analysed using Fiji. Raw Integrated density and cell perimeter were measured by manually following the contour of the cell using the segmented line tool (width=12 nm). Circularity (the ratio between longer and shortest axes for each cell) was determined by manually following the contour of the cell using the free hand line tool. The selected areas were then saved in the ROI manager for analysis. Single per unit area was calculated by (Raw Integrated density / area selected using the segmented line tool (width=12 nm) = single per unit area).

### Cell association assay of fluorescent monosaccharide polymers

U937 cells transfected with human DC-SIGN (U937-DC-SIGN) and controls U937 cells were obtained from the American Type Culture Collection (ATCC) and grown in suspension in RPMI-1640 medium (R0883, Sigma) containing 10% (v/v) foetal bovine serum (FBS, F9665, Sigma), 2 mM L-glutamine (35050-038, Gibco), 10 mM HEPES buffer (15630-056Gibco), 1 mM sodium pyruvate (11360-039, Gibco), 4.5 g/L D-glucose (A24940-01,Gibco) and 0.15 % (v/v) sodium bicarbonate (S8761, Sigma). U937 cells and moDCs were harvested by centrifugation at 250 x g for 5 min, washed in opti-MEM serum-free media (Gibco), re-suspended in opti-MEM, adjusted to 1.25 x10^6^ cells/ml and plated in 24 well tissue culture plates (250,000 cells/well, Costar) and incubated for 30 min at 37°C. Fluorescent monosaccharide polymers: Lewis^X^-PAA-FITC, fucose-PAA-FITC or galactose-PAA-FITC (Lectinity) were added to each well (5 μg/ml final) and opti-MEM to controls. Cultures were incubated for 1 h at 37°C and then transferred to fluorescein-activated cell sorting (FACS) tubes and 1 ml of FACS Buffer (0.5% (v/v) FBS, 15 mM NaN3 in PBS with Ca^2+^ and Mg^2+^ (D8537, Sigma)) was added to each tube. Cells were washed twice in FACS buffer by centrifugation at 300 x g for 5 min and re-suspended in 200μl of FACS buffer and fixed with 2% (v/v) Paraformaldehyde in PBS. Cells were analysed using Beckman Coulter FC500 flow cytometer. Data was analysed using Kaluza analysis software 1.5a. For cell association inhibition assays cells were pre-incubated with HMW, or opti-MEM for 1 h at 37°C before addition of fluorescent monosaccharide polymers. A similar procedure was used for CHO and CHO-MR cells but, being adherent cells, all washes were done in the wells and cells were harvested using trypsin-EDTA for analysis as described (45).

### Statistical analysis

Statistical analysis was performed in GraphPad Prism.

## Acknowledgements

This project was funded by The Medical Research Council project MR/P001033/1 to LMP, MC and PW. MA was funded by King Khalid University, Saudi Arabia. YA is funded by Shaqra University, Saudi Arabia, KL is funded by MRC-IMPACT PhD studentship. YI was funded by FP7 Marie Curie Fellowship grant PIIF-GA-2012-329832 and is receiving funding from the University of Tartu Institute of Technology. MC, LMP and PW are also funded by the National Biofilms Innovation Centre (NBIC) which is an Innovation and Knowledge Centre funded by the Biotechnology and Biological Sciences Research Council, Innovate UK and Hartree Centre [Award Number BB/R012415/1]. The funders had no role in study design, data collection and analysis, decision to publish, or preparation of the manuscript.

## Supporting Information

**Figure S1.** Point mutations identified within the *psl* operon in the P. aeruginosa wound isolates used in this study.

**Figure S2.** Generation of PAO1 biofilms under standardized flow conditions

**Figure S3.** Characterisation of HMW carbohydrate preparations

**Figure S4.** Binding of MR and DC-SIGN to HMW carbohydrate preparations is dose and Ca^2+^ dependent.

**Figure S5.** ^1^H-NMR analysis of HMW-1 and MHW-2.

**Figure S6.** Analysis of DC-SIGN and MR expression by moDCs

**Figure S7.** LPS-free HMW-2 does not affect cytokine production by moDCs in response LPS.

**Figure S8.** LPS-free HMW-2 does not influence the late response of moDCs to LPS.

**Figure S9.** LPS-free HMW-2 does not affect cytokine production by moDCs in response to Psl-deficient biofilms or planktonic cultures.

**Figure S10.** Specific detection of DC-SIGN in human DCs in the presence and absence of HMW-2.

**Figure S11.** Polymyxin B does not affect HMW-1 and HMW-2-mediated inhibition of Lewis^x^ uptake by moDCs.

**Figure S12.** Analysis of human MR binding to Lewis^x^-PAA using surface plasmon resonance.

**Figure S13.** LPS removal does not affect the ability of HMW-2 to inhibit ligand internalisation by DC-SIGN and MR expressing cells.

**Figure S14.** Biofilm carbohydrate composition does not influence cytokine production by moDCs.

**Video S1.** Binding of DC-SIGN to PAO1 biofilms

**Video S2.** Binding of MR (CTLD4-7) to PAO1 biofilms

**Video S3.** Secondary antibody control for DC-SIGN

**Video S4.** Secondary antibody control for MR (CTLD4-7)

**Video S5.** Binding of MR (CR-FNII-CTLD1-3) to PAO1 biofilms

## References

1. Mulcahy LR, Isabella VM, Lewis K. Pseudomonas aeruginosa biofilms in disease. Microb Ecol. 2014;68(1):1–12.

2. Davies JC, Bilton D. Bugs, biofilms, and resistance in cystic fibrosis. Respiratory care. 2009;54(5):628–40.

3. O’Sullivan BP, Freedman SD. Cystic fibrosis. Lancet. 2009;373(9678):1891–904.

4. Breidenstein EB, de la Fuente-Nunez C, Hancock RE. Pseudomonas aeruginosa: all roads lead to resistance. Trends Microbiol. 2011;19(8):419–26.

5. Jensen PO, Givskov M, Bjarnsholt T, Moser C. The immune system vs. Pseudomonas aeruginosa biofilms. FEMS Immunol Med Microbiol. 2010;59(3):292–305.

6. Colvin KM, Irie Y, Tart CS, Urbano R, Whitney JC, Ryder C, et al. The Pel and Psl polysaccharides provide Pseudomonas aeruginosa structural redundancy within the biofilm matrix. Environmental microbiology. 2012;14(8):1913–28.

7. Irie Y, Borlee BR, O’Connor JR, Hill PJ, Harwood CS, Wozniak DJ, et al. Self-produced exopolysaccharide is a signal that stimulates biofilm formation in Pseudomonas aeruginosa. Proceedings of the National Academy of Sciences of the United States of America. 2012;109(50):20632–6.

8. Franklin MJ, Nivens DE, Weadge JT, Howell PL. Biosynthesis of the Pseudomonas aeruginosa Extracellular Polysaccharides, Alginate, Pel, and Psl. Frontiers in microbiology. 2011;2:167.

9. Byrd MS, Sadovskaya I, Vinogradov E, Lu H, Sprinkle AB, Richardson SH, et al. Genetic and biochemical analyses of the Pseudomonas aeruginosa Psl exopolysaccharide reveal overlapping roles for polysaccharide synthesis enzymes in Psl and LPS production. Mol Microbiol. 2009;73(4):622–38.

10. Jennings LK, Storek KM, Ledvina HE, Coulon C, Marmont LS, Sadovskaya I, et al. Pel is a cationic exopolysaccharide that cross-links extracellular DNA in the Pseudomonas aeruginosa biofilm matrix. Proceedings of the National Academy of Sciences of the United States of America. 2015;112(36):11353–8.

11. Martinez-Pomares L. The mannose receptor. Journal of leukocyte biology. 2012;92(6):1177–86.

12. Garcia-Vallejo JJ, van Kooyk Y. The physiological role of DC-SIGN: a tale of mice and men. Trends Immunol. 2013;34(10):482–6.

13. Lam JS, Taylor VL, Islam ST, Hao Y, Kocincova D. Genetic and Functional Diversity of Pseudomonas aeruginosa Lipopolysaccharide. Frontiers in microbiology. 2011;2:118.

14. Linehan SA, Martinez-Pomares L, da Silva RP, Gordon S. Endogenous ligands of carbohydrate recognition domains of the mannose receptor in murine macrophages, endothelial cells and secretory cells; potential relevance to inflammation and immunity. Eur J Immunol. 2001;31(6):1857–66.

15. Martinez-Pomares L, Wienke D, Stillion R, McKenzie EJ, Arnold JN, Harris J, et al. Carbohydrate-independent recognition of collagens by the macrophage mannose receptor. Eur J Immunol. 2006;36(5):1074–82.

16. Irie Y, Starkey M, Edwards AN, Wozniak DJ, Romeo T, Parsek MR. Pseudomonas aeruginosa biofilm matrix polysaccharide Psl is regulated transcriptionally by RpoS and post-transcriptionally by RsmA. Mol Microbiol. 2010;78(1):158–72.

17. Pestrak MJ, Chaney SB, Eggleston HC, Dellos-Nolan S, Dixit S, Mathew-Steiner SS, et al. Pseudomonas aeruginosa rugose small-colony variants evade host clearance, are hyper-inflammatory, and persist in multiple host environments. PLoS Pathog. 2018;14(2):e1006842.

18. Murphy K, Park AJ, Hao Y, Brewer D, Lam JS, Khursigara CM. Influence of O polysaccharides on biofilm development and outer membrane vesicle biogenesis in Pseudomonas aeruginosa PAO1. J Bacteriol. 2014;196(7):1306–17.

19. Bales PM, Renke EM, May SL, Shen Y, Nelson DC. Purification and Characterization of Biofilm-Associated EPS Exopolysaccharides from ESKAPE Organisms and Other Pathogens. PloS one. 2013;8(6):e67950.

20. Lowman DW, West LJ, Bearden DW, Wempe MF, Power TD, Ensley HE, et al. New insights into the structure of (1-->3,1-->6)-beta-D-glucan side chains in the Candida glabrata cell wall. PloS one. 2011;6(11):e27614.

21. Zhao K, Tseng BS, Beckerman B, Jin F, Gibiansky ML, Harrison JJ, et al. Psl trails guide exploration and microcolony formation in Pseudomonas aeruginosa biofilms. Nature. 2013;497(7449):388–91.

22. Feinberg H, Guo Y, Mitchell DA, Drickamer K, Weis WI. Extended neck regions stabilize tetramers of the receptors DC-SIGN and DC-SIGNR. The Journal of biological chemistry. 2005;280(2):1327–35.

23. Relloso M, Puig-Kroger A, Pello OM, Rodriguez-Fernandez JL, de la Rosa G, Longo N, et al. DC-SIGN (CD209) expression is IL-4 dependent and is negatively regulated by IFN, TGF-beta, and anti-inflammatory agents. Journal of immunology. 2002;168(6):2634–43.

24. Liu Y, Chirino AJ, Misulovin Z, Leteux C, Feizi T, Nussenzweig MC, et al. Crystal structure of the cysteine-rich domain of mannose receptor complexed with a sulfated carbohydrate ligand. J Exp Med. 2000;191(7):1105–16.

25. Yamada KJ, Kielian T. Biofilm-Leukocyte Cross-Talk: Impact on Immune Polarization and Immunometabolism. J Innate Immun. 2019;11(3):280–8.

26. Geijtenbeek TB, van Kooyk Y. Pathogens target DC-SIGN to influence their fate DC-SIGN functions as a pathogen receptor with broad specificity. APMIS. 2003;111(7-8):698–714.

27. Nigou J, Zelle-Rieser C, Gilleron M, Thurnher M, Puzo G. Mannosylated lipoarabinomannans inhibit IL-12 production by human dendritic cells: evidence for a negative signal delivered through the mannose receptor. Journal of immunology. 2001;166(12):7477–85.

28. Borlee BR, Goldman AD, Murakami K, Samudrala R, Wozniak DJ, Parsek MR. Pseudomonas aeruginosa uses a cyclic-di-GMP-regulated adhesin to reinforce the biofilm extracellular matrix. Mol Microbiol. 2010;75(4):827–42.

29. Xaplanteri P, Lagoumintzis G, Dimitracopoulos G, Paliogianni F. Synergistic regulation of Pseudomonas aeruginosa-induced cytokine production in human monocytes by mannose receptor and TLR2. Eur J Immunol. 2009;39(3):730–40.

30. Vendele I, Willment JA, Silva LM, Palma AS, Chai W, Liu Y, et al. Mannan detecting C-type lectin receptor probes recognise immune epitopes with diverse chemical, spatial and phylogenetic heterogeneity in fungal cell walls. PLoS Pathog. 2020;16(1):e1007927.

31. Passos da Silva D, Matwichuk ML, Townsend DO, Reichhardt C, Lamba D, Wozniak DJ, et al. The Pseudomonas aeruginosa lectin LecB binds to the exopolysaccharide Psl and stabilizes the biofilm matrix. Nat Commun. 2019;10(1):2183.

32. Li H, Mo K-F, Wang Q, Stover CK, DiGiandomenico A, Boons G-J. Epitope Mapping of Monoclonal Antibodies using Synthetic Oligosaccharides Uncovers Novel Aspects of Immune Recognition of the Psl Exopolysaccharide of Pseudomonas aeruginosa. Chemistry – A European Journal. 2013;19(51):17425–31.

33. Ray VA, Hill PJ, Stover CK, Roy S, Sen CK, Yu L, et al. Anti-Psl Targeting of Pseudomonas aeruginosa Biofilms for Neutrophil-Mediated Disruption. Scientific Reports. 2017;7(1):16065.

34. Lowman DW, Ensley HE, Greene RR, Knagge KJ, Williams DL, Kruppa MD. Mannan structural complexity is decreased when Candida albicans is cultivated in blood or serum at physiological temperature. Carbohydr Res. 2011;346(17):2752–9.

35. McCarthy RR, Mazon-Moya MJ, Moscoso JA, Hao Y, Lam JS, Bordi C, et al. Cyclic-di-GMP regulates lipopolysaccharide modification and contributes to Pseudomonas aeruginosa immune evasion. Nat Microbiol. 2017;2:17027.

36. Geijtenbeek TB, Torensma R, van Vliet SJ, van Duijnhoven GC, Adema GJ, van Kooyk Y, et al. Identification of DC-SIGN, a novel dendritic cell-specific ICAM-3 receptor that supports primary immune responses. Cell. 2000;100(5):575–85.

37. Geijtenbeek TB, Krooshoop DJ, Bleijs DA, van Vliet SJ, van Duijnhoven GC, Grabovsky V, et al. DC-SIGN-ICAM-2 interaction mediates dendritic cell trafficking. Nat Immunol. 2000;1(4):353–7.

38. Karosi T, Csomor P, Hegyi Z, Sziklai I. The presence of CD209 expressing dendritic cells correlates with biofilm positivity in chronic rhinosinusitis with nasal polyposis. Eur Arch Otorhinolaryngol. 2013;270(9):2455–63.

39. Ochoa MT, Loncaric A, Krutzik SR, Becker TC, Modlin RL. “Dermal dendritic cells” comprise two distinct populations: CD1+ dendritic cells and CD209+ macrophages. The Journal of investigative dermatology. 2008;128(9):2225–31.

40. Murphy BS, Bush HM, Sundareshan V, Davis C, Hagadone J, Cory TJ, et al. Characterization of macrophage activation states in patients with cystic fibrosis. Journal of cystic fibrosis: official journal of the European Cystic Fibrosis Society. 2010;9(5):314–22.

41. Tailleux L, Pham-Thi N, Bergeron-Lafaurie A, Herrmann JL, Charles P, Schwartz O, et al. DC-SIGN induction in alveolar macrophages defines privileged target host cells for mycobacteria in patients with tuberculosis. PLoS Med. 2005;2(12):e381.

42. Anthony RM, Kobayashi T, Wermeling F, Ravetch JV. Intravenous gammaglobulin suppresses inflammation through a novel T(H)2 pathway. Nature. 2011;475(7354):110–3.

43. Schaefer M, Reiling N, Fessler C, Stephani J, Taniuchi I, Hatam F, et al. Decreased pathology and prolonged survival of human DC-SIGN transgenic mice during mycobacterial infection. Journal of immunology. 2008;180(10):6836–45.

44. Schindelin J, Arganda-Carreras I, Frise E, Kaynig V, Longair M, Pietzsch T, et al. Fiji: an open-source platform for biological-image analysis. Nat Methods. 2012;9(7):676–82.

45. Su Y, Bakker T, Harris J, Tsang C, Brown GD, Wormald MR, et al. Glycosylation influences the lectin activities of the macrophage mannose receptor. The Journal of biological chemistry. 2005;280(38):32811–20.

46. Holloway BW, Krishnapillai V, Morgan AF. Chromosomal Genetics of Pseudomonas. Microbiol Rev. 1979;43(1):73–102.

47. Hickman JW, Tifrea DF, Harwood CS. A chemosensory system that regulates biofilm formation through modulation of cyclic diguanylate levels. P Natl Acad Sci USA. 2005;102(40):14422–7.

